# PANINI: Pangenome Neighbor Identification for Bacterial Populations

**DOI:** 10.1101/174409

**Authors:** Khalil Abudahab, Joaquín M. Prada, Zhirong Yang, Stephen D. Bentley, Nicholas J. Croucher, Jukka Corander, David M. Aanensen

## Abstract

The standard workhorse for genomic analysis of the evolution of bacterial populations is phylogenetic modelling of mutations in the core genome. However, in the current era of population genomics, a notable amount of information about evolutionary and transmission processes in diverse populations can be lost unless the accessory genome is also taken into consideration. Here we introduce PANINI, a computationally scalable method for identifying the neighbours for each isolate in a data set using unsupervised machine learning with stochastic neighbour embedding. PANINI is browser-based and integrates with the Microreact platform for rapid online visualisation and exploration of both core and accessory genome evolutionary signals together with relevant epidemiological, geographic, temporal and other metadata. Several case studies with single-and multi-clone pneumococcal populations are presented to demonstrate ability to identify biologically important signals from gene content data. PANINI is available at <u>http://panini.wgsa.net/</u> and code at <u>http://gitlab.com/cgps/panini</u>

## BACKGROUND

In less than a decade, bacterial population genomics has progressed from sequencing of dozens to thousands of strains [1,2,3,4]. The biological insights enabled by population genomics are particularly important in evolutionary epidemiology, as the genome sequences provide high resolution data for the estimation of transmission and evolutionary dynamics, including horizontal transfer of virulence and resistance elements. Phylogenetic trees are the main framework utilised for visualisation and exploration of population genomic data, both in terms of the level of relatedness of strains and for mapping relevant metadata such as geographic locations and host characteristics [5]. While trees are highly useful, they are in general estimated using only core genome variation (i.e. those regions of the genome common to all members of a sample), which may represent only a fraction of the relevant differences present in genomes across the study population. Several recent studies highlight the importance of considering variation in gene content when investigating the ecological and evolutionary processes leading to the observed data [6, 7].

The rapidly increasing size of population genomic datasets calls for efficient visualisation methods to explore patterns of relatedness based on core genomic polymorphisms, accessory gene content, epidemiological, geographical and other metadata. Here we introduce a framework that integrates within the web application Microreact [5], by utilising a popular unsupervised machine learning technique for big data to infer neighbors of bacterial strains from accessory gene content data and to efficiently visualize the resulting relationships. The machine learning method, called t-SNE, has already gained widespread popularity for exploring image, video and textual data [8,9], but has to our knowledge not yet been utilized for bacterial population genomics.

Since gene content may in general be rapidly altered in bacteria, it provides a high-resolution evolutionary marker of relatedness which can extend far beyond core genome mutations [7]. Different processes driving horizontal movement of DNA, such homologous recombination, conjugative transfer of plasmids and phage infections, all affect the gene content within and outside of a chromosome. By contrasting core and non-core gene content, one can investigate and draw conclusions about genome dynamics across a sample collection. Here we demonstrate the biological utility of such an approach by application to multiple population data sets.

## METHODS AND RESULTS

Student t-distributed Stochastic Neighbor Embedding (t-SNE) is a machine learning algorithm which is widely used for data visualization [8]. It is suitable for embedding a set of high-dimensional data items in a 2-or 3-dimensional space. The embedding approximately preserves the pairwise similarities between the data items.

The t-SNE algorithm consists of two main steps. First, it calculates the similarities between the data items in the high-dimensional space, which is typically based on normal distribution around each data item. The similarities are then normalized to be probabilities (i.e. they sum to one). Similarities in the low-dimensional space are analogously defined and normalized except that Student t-distribution replaces the Gaussians. Second, t-SNE minimizes Kullback-Leibler divergence between the two probability matrices over the embedding coordinates. Finally, the 2-D t-SNE result can be visualized as a scatter plot where each dot indicates a data item.

t-SNE as an unsupervised method is particularly useful for exploratory data analysis. It has a wide range of applications in music analysis, cancer research, computer security research, bioinformatics, and biomedical signal processing. In many cases, t-SNE is able to identify meaningful data structures such as clusters even without feature engineering or structural assumptions, e.g. about number of clusters underlying the data. Here, we use the latest version of the t-SNE projection method, adopting the Barnes-Hut algorithm for accelerating the divergence minimization [9].

To demonstrate utility within population genomics, firstly, we explore how the method performs in a simulated setting, where the relationship between all sequences is known and then extended our analysis to published bacterial population data sets, allowing us to uncover previously unseen relationships between data and to address important biological questions.

## SIMULATED DATA

To our knowledge, this is the first time that the t-SNE projection method has been used to explore patterns of genetic relatedness between different bacterial isolates. We have therefore validated the methodology by assessing how well it identifies neighbours and clusters for simulated genetic sequences. Firstly, we randomly generated multiple synthetic datasets of related isolates, with each defined as a sequence of present/absent genes. Each dataset is generated using the following parameters:

1. There are 20 clusters as underlying subpopulations.
2. The number of isolates belonging to a cluster is drawn from a Poisson distribution with mean 15.
3. Each cluster is defined by a number of core genes, which ranges uniformly from 1 to 100.
4. Each isolate has a probability between 80% to 99% of independently carrying each of the core genes of the cluster it belongs to.
5. Conversely, each isolate has a probability (PN) to independently carry each of the non-core genes of its cluster. Non-core genes are composed of core genes of other clusters and "noise" genes which are not defining characteristics of any cluster (in total 300 genes).

Each generated dataset has on average 300 isolates with a gene content of 1300 genes present/absent on average. For each dataset, we estimated the genetic pairwise Hamming distance (d_H_) and the distance using the t-SNE algorithm (d_t_). The Hamming distance is simply the number of differences between two sequences of equal length, which in this case refers to a gene being present in one isolate but absent in the other. The implementation of the t-SNE algorithm that we use yields a coordinate in a 2D plane for each isolate, and we calculate the distance d_t_ simply as the Euclidean distance for each pair of isolates.

If a cluster is sufficiently differentiable in terms of its gene content, we expect the Hamming distance within the cluster to be smaller than to any other isolate not belonging to it. For the t-SNE algorithm to be considered valid, it should be able to project the isolates from the same cluster on the 2D plane sufficiently close together so that the Euclidean distance within the cluster is smaller than to any other isolate. Given the conditions that were used to generate the synthetic datasets, not all clusters are necessarily differentiable in terms of their gene content, therefore we classified the t-SNE algorithm as performing erroneously only when a pair of isolates belonging to different cluster are not identified as such by the algorithm but are correctly identified using the Hamming distance. For high levels of noise, i.e. a large value of PN, differentiating the clusters using their gene content becomes increasingly difficult as the isolates may lack a sufficiently stable signal of relatedness.

We analyzed the performance of the t-SNE algorithm for three levels of noise PN: 0.001, 0.005 and 0.01, which measures the average proportion of non-core genes in each isolate. We performed 100 repeats for each noise value, which for each repeat involves generating on average 300 sequences and comparing almost 45000 pairs of isolates. The average error for the three noise values was 0.5%, 1% and 4% respectively, with a small error representing a particular isolate mis-allocated (i.e. very close to a different cluster) and a large error representing two clusters which are not appropriately differentiated by the t-SNE algorithm, illustrated in Figure 1. The error of the t-SNE algorithm increases with the noise, as shown in Figure 1(iii), and with the total number of clusters (not shown).

**Figure 1.**
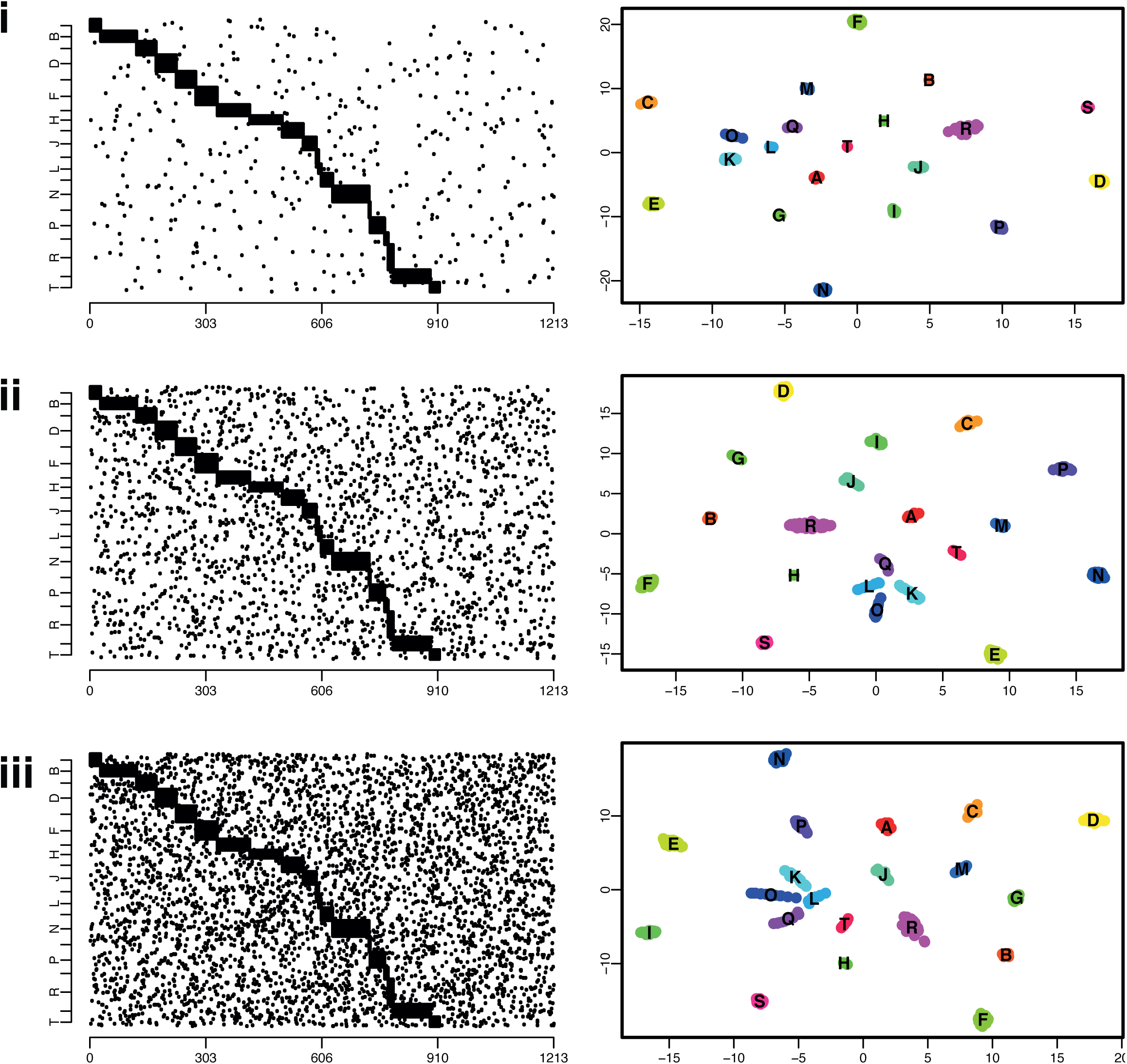
Illustration of a simulated dataset, with the isolates’ gene content (left), black dots indicate presence of a gene, x-axis represent all the considered genes (total of 1213 genes in this simulation). The right panels show the embedded locations in the 2D plane as estimated by the t-SNE algorithm, with each colour representing a cluster in the underlying simulation model. Clusters are named using the alphabet (A, B, C…). From top to bottom, plots indicate simulations generated with 0.1% (i), 0.5% (ii) and 1%(iii) noise, respectively.

## WEB APPLICATION - <u>https://panini.wgsa.net/</u>

The t-SNE algorithm implemented in C++ (<u>https://github.com/lvdmaaten/</u> <u>bhtsne</u>) was wrapped as a Node.js native module and embedded within a web application. The application is written in JavaScript and utilises React (<u>https://facebook.github.io/react</u>) for front-end and the Vis.js library (<u>http://visjs.org</u>) for network visualisation.

1) Data are uploaded as a gene presence/absence matrix - Panini expects data in the .rtab format (the output from Roary: the pan genome pipeline [10]; <u>https://sanger-pathogens.github.io/Roary</u>) However, this is simply a data file containing gene rows and isolate columns with ‘1’ or ‘0’ indicating presence/absence of a particular gene for a particular isolate.

2) Genes present in all isolates are ignored (i.e. core genome) and non-core genes are clustered using t-SNE with default parameters (auto perplexity and theta=0.5 – parameters can be changed by users).

3) The Results (x, y coordinates, a ‘.dot’ format file containing graph layout, csv and JSON) are made available for download and reuse. Results are also visualised directly within the PANINI web application as a graph layout.

To interpret the data in an epidemiological, phylogeographic and geographic context, the estimated network can also be uploaded directly to the Microreact platform allowing a user to add other forms of data to relate to the resulting neighbor embedding, typically a phylogenetic tree, geographical locations of the isolates, and temporal data (Further information and instructions at <u>https://microreact.org</u>).

## UTILITY WITH EXISTING PUBLISHED DATASETS

To demonstrate utility of t-SNE clustering we applied the method to three published datasets which used Whole Genome Sequencing (WGS) to study the evolution of the bacterium *Streptococcus pneumoniae*. The first, a population level dataset, detailed population-wide diversity of pneumococci within Massachusetts, USA pre- and post vaccine introduction [2], while the second and third detailed international collections of globally-disseminated multidrug-resistant lineages of *Streptococcus pneumoniae* [11, 12]. Additional biological insights made possible with PANINI are described, and links to the projects within Microreact for further exploration of associated metadata and download of raw data formats are provided.

## ANALYSIS OF A DIVERSE PNEUMOCOCCAL POPULATION

### Data Visualisation and download

<u>https://microreact.org/project/panini-sparc?ui=nt</u>

### Source data and .RTab file

<u>https://gitlab.com/cgps/panini/datasets/tree/master/SPARC</u>

### Video walkthrough for PANINI and Microreact creation/use: <u>https://vimeo.com/230416235</u>

PANINI was applied to a collection of 616 systematically-sampled pneumococcal isolates from a vaccine and antimicrobial resistance surveillance project in Massachusetts [13]. The original analysis of gene content in this collection identified 5,442 ‘clusters of orthologous genes’ (COGs) [2], the core set of which was used to define fifteen ‘sequence clusters’ with BAPS (<u>http://www.helsinki.fi/bsg/software/BAPS</u>) [18]. For most of the sequence clusters, the correspondence between a group in the PANINI output and the original sequence clusters was exact (Figure 2A), reflecting their similarity both in terms of the core and accessory genomes [14]. These sets of isolates therefore represent well-defined, distinct lineages. However, SC1, SC6, SC10 and SC12 all exhibited distinct substructuring in the PANINI output. This corresponded well with the diverse core genome observed in these clusters (Figure 2B), and in each case, these groups were consistent with clades within the sequence clusters. These sequence clusters are therefore likely to represent amalgams of genotypes that should be subdivided into multiple clusters. Conversely, PANINI revealed clear substructuring within the previously unclustered SC16, which was also consistent with the core genome phylogeny. Hence PANINI can easily facilitate the division of a diverse population into discrete genotypes that are coherent in their accessory and core genome content.

**Figure 2.**
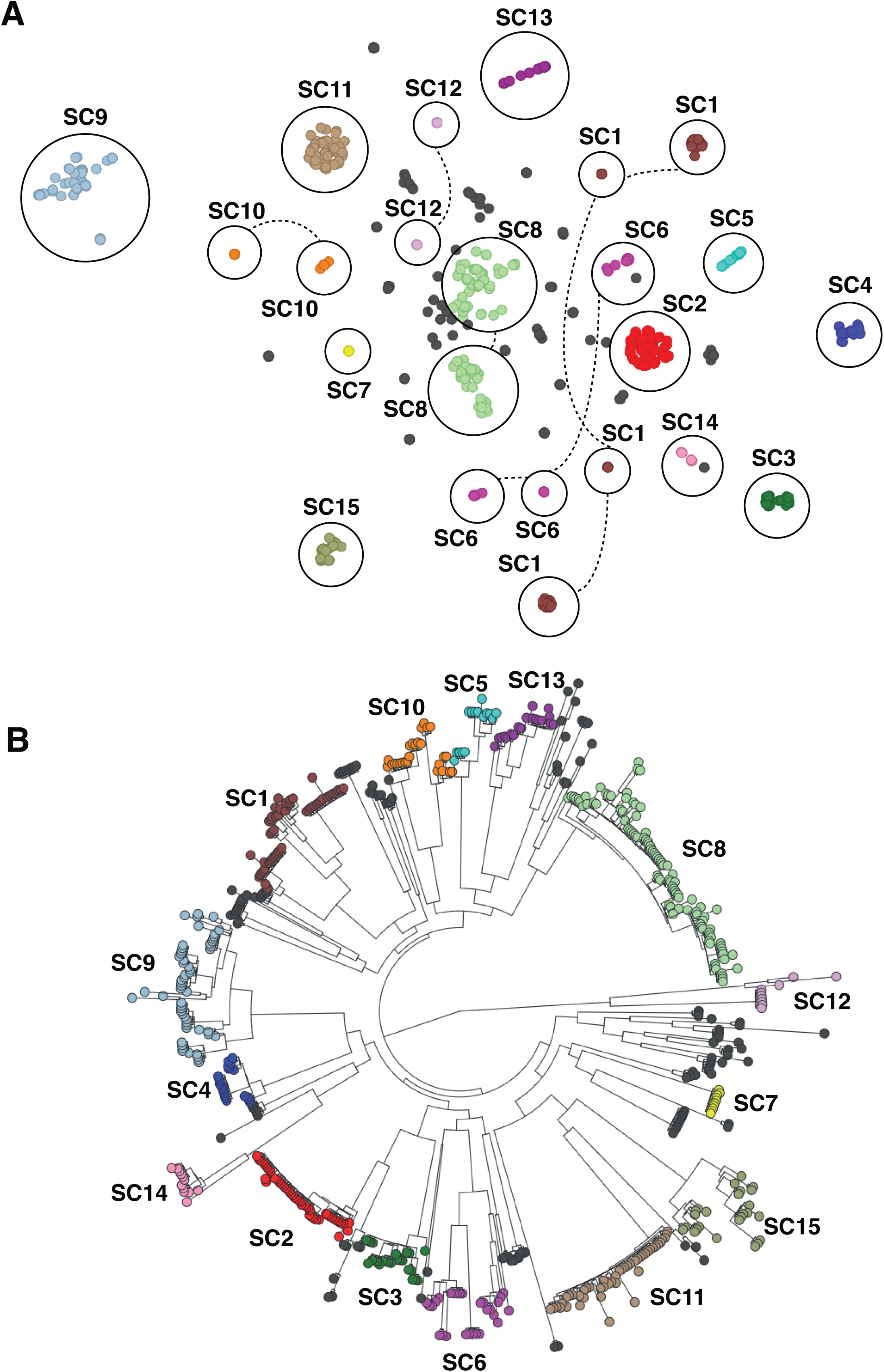
A) Annotated output of the PANINI algorithm applied to 616 *S. pneumoniae* isolates from a diverse population in Massachusetts. Each node represents an isolate, each of which is coloured according to its sequence cluster, as defined using the core genome. Clusters of isolates belonging to the same sequence cluster are circled and annotated. Where sequence clusters are divided into multiple groups in the PANINI network, the circles are joined by dashed lines. B) Core genome phylogeny based on comparison of conserved clusters of orthologous genes adapted from [2] and displayed within Microreact. Sequence clusters are annotated for comparison with non-core clustering.

## EXTENSIVE PROPHAGE VARIATION IN A MULTIDRUG-RESISTANT LINEAGE

### Data Visualisation and download: <u>https://microreact.org/project/panini-pmen2?ui=nt</u>

### Source data and .RTab file

<u>https://gitlab.com/cgps/panini/datasets/tree/master/PMEN2</u>

PANINI was applied to an analysis of orthologous genes across a global collection of 190 isolates from the multidrug-resistant *Streptococcus pneumoniae* clone PMEN2 [11], which caused a large outbreak of disease in Iceland starting in the late 1980s (Figure 3A). Multiple distinct clusters were again evident in the output (Figure 3B). In some cases, these were consistent with the phylogeny. The original analysis identified two independent entries of the lineage into Iceland, clades IC1 and IC2, the latter of which contained many fewer isolates and was clustered as IcA in the annotated output. By contrast, IC1 was distributed across four clusters IcB-E, which did not correspond with clear clades in the phylogeny. The difference between IcB and IcC is technical, rather than biological: all IcB isolates were sequenced early in the project with 54 nt reads, whereas most IcC isolates were sequenced with 75 nt reads. Unusually for pneumococci, the isolates in both these groups were trilysogenic, carrying prophage similar to ϕ670-6B.1 and ϕ670-6B.2, found in the *S. pneumoniae* 670-6B genome inserted between *dnaN* and *pth* (*att*_670_), and within the *comYC* gene (*att*_comYC_), respectively; and a prophage isolated from 0211+13275, inserted at SPN23F15280 - SPN23F15810 (*att*_MM1_) [15]. The apparent rapid acquisition, and stable maintenance, of multiple viral loci may relate to the abrogation of these bacteria’s competence system by the insertion of prophage ϕIC1 into *comYC* [11,16]. Group IcD, interspersed with IcB and IcC within clade IC1 in the phylogeny, differs in the absence of prophage similar to ϕ670-6B.2. IcE, also polyphyletic within clade IC1, differed in having lost the region of PPI-1 that encodes the *pia* iron transport operon, which plays a role in pneumococcal pathogenesis in animal models [17]. Hence it is not surprising to find these isolates were only recovered from sputum, otitis media samples or nasopharyngeal swabs.

**Figure 3.**
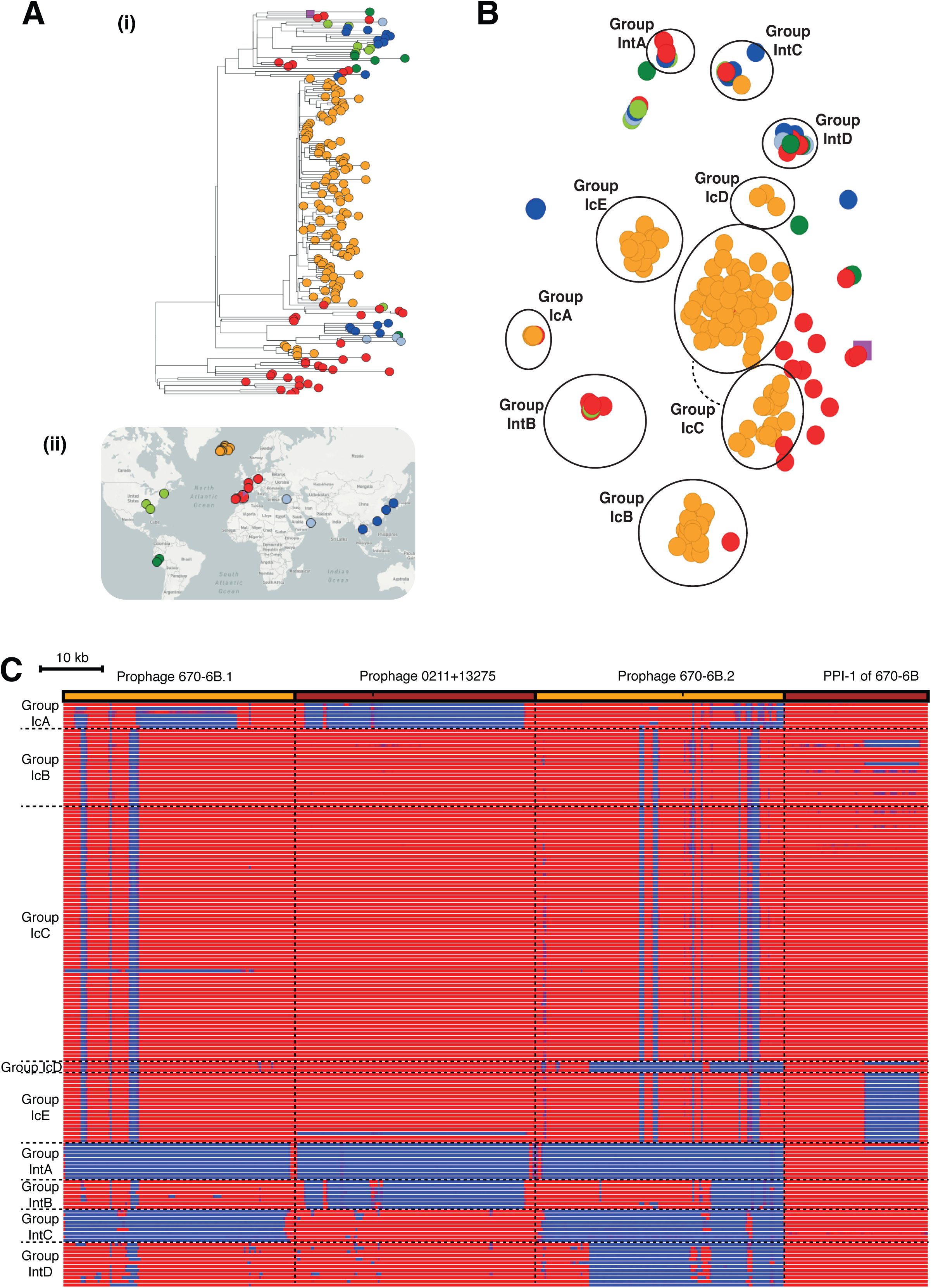
Analysis of the *S. pneumoniae* PMEN2 lineage. A) i) Core genome phylogeny with tree leaves coloured by country of origin and ii) geographic origin of isolates. B) Annotated output of the PANINI algorithm applied to 189 isolates from an international collection of representatives of the *S. pneumoniae* PMEN2 lineage. Each point is coloured according to its region of origin. Groups defined by the structure of the PANINI output are circled and annotated. Clusters containing primarily Icelandic isolates (coloured orange) are labelled with ‘Ic’ prefixes, whereas those containing isolates from multiple countries are labelled with ‘Int’ prefixes. C) Variation in accessory loci associated with differential classification of isolates into groups. The orange and brown bands across the top of the figure indicate the extent of the three prophage and pneumococcal pathogenicity island 1 (PPI-1) sequences, against which the short read data from the isolates were mapped. The heatmap below includes one row per isolate, which were ordered according to their grouping in panel A. The heatmap is coloured blue where mapping coverage was low, indicating a locus is absent, and red were mapping coverage was high, indicating a sequence was present. Horizontal dashed lines indicate the boundaries between the groups of isolates, which vertical dashed lines indicate the boundaries between loci.

Multiple distinct clusters of non-Icelandic isolates were also observed. These all represented cases where t-SNE grouped isolates that were disparate in terms of their country and year of isolation, as well as having a polyphyletic distribution across the whole genome phylogeny. These groupings represented cases of convergent evolution through parallel acquisition very similar prophage. Group IntA lacked any prophage similar to those shown in Figure 3C; group IntB had prophage with some similarity to both prophage in the reference genome; group IntC only had a prophage with similarity to ϕ0211+13275, whereas group IntD had prophage similar to ϕ0211+13275 and ϕ670-6B.1 as well. Hence the rapid movement of prophage sequences within lineages [14] clearly substantially contributes to the changes in gene content observed over short timescales. PANINI facilitates rapid analysis of these diverse elements, and their complex relationship with bacterial population structure.

## MOBILE ELEMENT AND SEROTYPE VARIATION IN A VACCINE-ESCAPE LINEAGE

### Data Visualisation and download

<u>https://microreact.org/project/panini-pmen14?ui=nt</u>

### Source data and .RTab file

<u>https://gitlab.com/cgps/panini/datasets/tree/master/PMEN14</u>

PANINI was similarly applied to 176 isolates of the multidrug-resistant *S. pneumoniae* PMEN14 lineage [11]. Although the sequences came from many countries, the collection was strongly enriched for bacteria from the Maela refugee camp in Thailand, which fell into five clades (ML1-5), of which ML2 was the largest. The groups identified by PANINI were again polyphyletic (Figure 4A), with ML2 split up in a similar manner to the PMEN2 clade IC1. This was again driven by the distribution of prophage sequence: group 1 isolates were free of prophage, whereas group 2 isolates were infected with a ‘group 2-type’ prophage, and group 3 isolates were infected with a similar, but distinct, ‘group 3-type’ prophage (Figure 4B). Clade ML2 isolates in group 4 were distinguished by variation in another mobile genetic element, a phage-related chromosomal island (PRCI), shared by most of the isolates. This PRCI was absent from these assemblies, either because at least part of the element had been lost through deletion, replacement with a related sequence (isolate 6259_1-15), or the acquisition of a second, highly similar PRCI that prevented effective assembly of either (isolates 6237_8-12, 6237_8-13 and 6237_8-18). In this latter case, mapping to the element was still evident.

**Figure 4.**
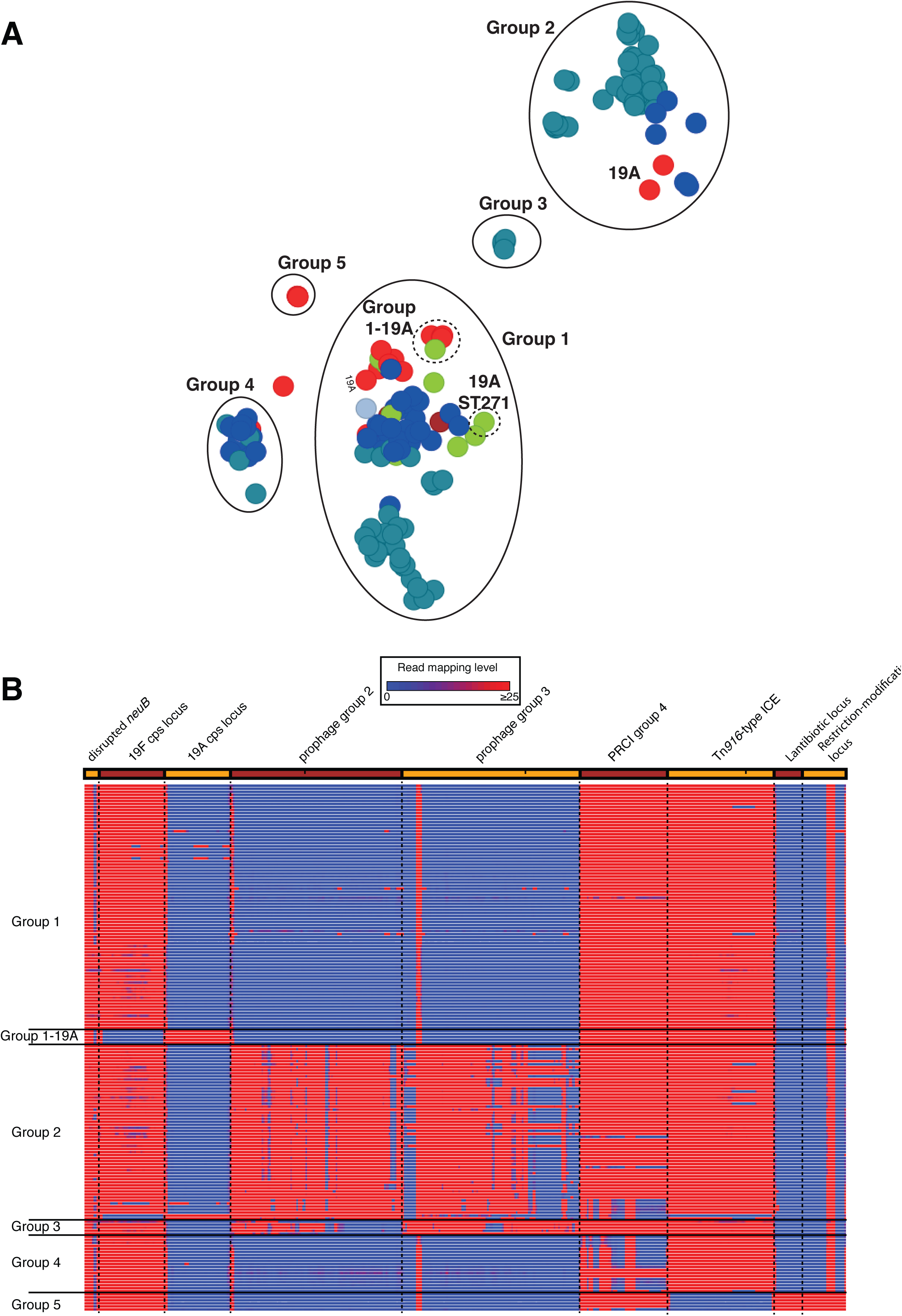
Analysis of the *S. pneumoniae* PMEN14 lineage. A) Annotated output of the PANINI algorithm applied to 176 isolates from an international collection of representatives of the *S. pneumoniae* PMEN14 lineage. The main groups 1-5 are circled with solid lines and named; the subgroups within group 1 are circled by dashed lines. (B) Variation in accessory loci associated with differential classification of isolates into groups. This heatmap is displayed as in Figure 3. In this case, the sequence loci across the top are more functionally diverse. The first is the *neuB* coding sequence with an IS*Spn*8 element inserted into it. The lack of mapping to the middle of this column indicates the absence of this insertion sequence anywhere in the chromosome. The next loci are alternative alleles of the capsule polysaccharide synthesis locus, one encoding for the biosynthesis of the 7-valent polysaccharide conjugate vaccine (PCV7) type 19F polysaccharide, the other for the non-PCV7 type 19A polysaccharide. These are followed by two similar prophage, one associated with group 2 isolates, the other with group 3 isolates; the similarity between these two viruses means there is extensive mapping to both, even when an isolate only contains one of them. The PRCI absent from the assemblies of group 4 isolates is next; mapping suggests this is actually present in some, but PANINI nevertheless included them in this group because the acquisition of a further, related PRCI prevented either assembling accurately. This is followed by the Tn*916* conjugative element, absent from the group 5 isolates, which possess genomic islands encoding for the biosynthesis of a lantibiotic and a restriction-modification system, included at the right-hand end of the panel.

A fifth group, which did not include any Maela isolates, corresponded to the antibiotic-susceptible outgroup isolates. These differed through the absence of a third type of mobile element, the Tn*916* integrative and conjugative element, an antibiotic resistance-encoding genomic island that was absent from these ‘outgroup’ isolates. Additionally, these bacteria shared two smaller genomic islands, encoding putative lantibiotic biosynthesis and restriction-modification operons, which were absent from the multidrug-resistant isolates. Variation in other non-mobile element islands was also detectable. The group 1-19A subcluster contained isolates of serotype 19A, produced through two independent serotype switching recombinations at the capsule polysaccharide synthesis (*cps*) locus that resulted in genotypes ‘19A ST320’ and ‘19A ST236’. These changes were responsible for allowing isolates to evade the seven valent polysaccharide conjugate vaccine, which targeted the lineage’s ancestral serotype 19F, expressed by almost all the rest of the collection [12]. A smaller serotype switching recombination, which did not replace the entire serotype-determining *cps* locus, generated the ‘19A ST271’ isolates [12]. The smaller associated change in gene content meant this isolate was not clearly distinguished from the rest of group 1 (Figure 4A).

## DISCUSSION

The rapid increase in sampling density of bacterial populations for epidemiological and evolutionary studies highlights the need of combining traditional genomic markers, such as SNP loci and small insertions or deletions in coding regions, with measures of difference in terms of gene content. As many bacteria have varied accessory genomes, changes in the gene content can offer a way to identify epidemiologically or evolutionarily important clues about the evolutionary processes affecting a pathogen’s spread. As we illustrated here, such information is most useful when clustering is combined within a phylogeographical approach, and visualized jointly in a seamless fashion enabling the rapid interpretation of core and non-core clustering in the context of where and when data were collected.

The t-SNE algorithm is a very efficient approach to cluster isolates based on their gene content. In the simulated scenarios considering synthetic data, the errors in clustering always remained small, either representing an isolate allocated to a wrong cluster, or two clusters which were not appropriately differentiated. However, this only occurred in simulations with the "noise" level much higher than expected in nature. In general, what we defined as "core" genes in a cluster rarely appear in isolates not belonging to the cluster, and if they do, it is typically at much lower frequencies than those we considered. Furthermore, in our synthetic datasets we formed clusters defined by as few as a single core gene. These clusters with a limited number of core genes, combined with relatively high levels of "noise", are in practice almost completely indistinguishable from others, as illustrated in Figure 1 (iii - clusters K, L, O and Q). Overall, our simulated datasets are conservative, as the gene absence and presence variation is higher than expected in natural populations, and therefore indicate that the t- SNE is a promising approach for rapidly and accurately clustering bacteria based on gene content.

When applied to a population-wide genomic dataset, the algorithm was clearly able to identify distinct lineages within a diverse collection. This analysis could highlight which clusters, defined using the core genome, could be sensibly subdivided, and which small groups of within a diverse set of strains could be justifiably regarded as new clusters. Within lineages, the same congruence between core and accessory genomes across clades was not observed. Instead, clusters were distinguished by rapidly occurring, homoplasic alterations, such as phage infection. In this context, PANINI provides an intuitive way in which to understand the distribution of rapidly-evolving aspects of the genome, which are difficult to analyse in a conventional phylogenetic framework. PANINI is therefore a promising platform through which biologically-important changes in bacterial gene content can be uncovered at all levels of evolutionary, ecological and epidemiological analyses.

